# The Tissue Engineering Grail: Seamless Biofabrication of Scaffold-free Hollow Constructs

**DOI:** 10.1101/2025.11.04.685834

**Authors:** Alireza Shahin-Shamsabadi, John Cappuccitti

**Affiliations:** Evolved.Bio, 280 Joseph Street, Kitchener, Ontario, Canada

**Keywords:** Scaffold-free tissue engineering, Anchored cell sheet engineering, Hollow tubes and spheres, Modular tissue engineering

## Abstract

Scaffold-free tissue engineering enables the construction of biomimetic tissues and organs by preserving cell-cell and cell-matrix interactions while avoiding exogenous scaffolds and biomaterials. Yet current approaches are limited to thin sheets or simple spheroids and often lack cellular maturity and organized extracellular matrix (ECM). Here, Anchored Cell Sheet Engineering, a concept that previously introduced anchors to guide the remodeling of cell sheets into more mature fibers or sheets, is extended to achieve seamless, single-step biofabrication of scaffold-free hollow tubular and spherical constructs for sustained biological and mechanical functions under physiological conditions. Using custom culture devices with curved geometries for two-dimensional (2D) culture, continuous confluent cell-ECM layers were formed that were then delaminated and guided by strategically positioned central cores with different shapes and sizes to undergo tension-mediated remodeling into mechanically stable hollow structures. This approach allows modulation of wall thickness, supports multi-layered architectures, and yields constructs capable of withstanding fluid flow. By expanding scaffold-free biofabrication beyond sheets and fibers to robust hollow geometries, this work establishes a versatile set of physiologically relevant building blocks for scalable bottom-up assembly of complex, multi-tissue organ-like constructs within a bioassembloid framework.

**Graphic Abstract:** 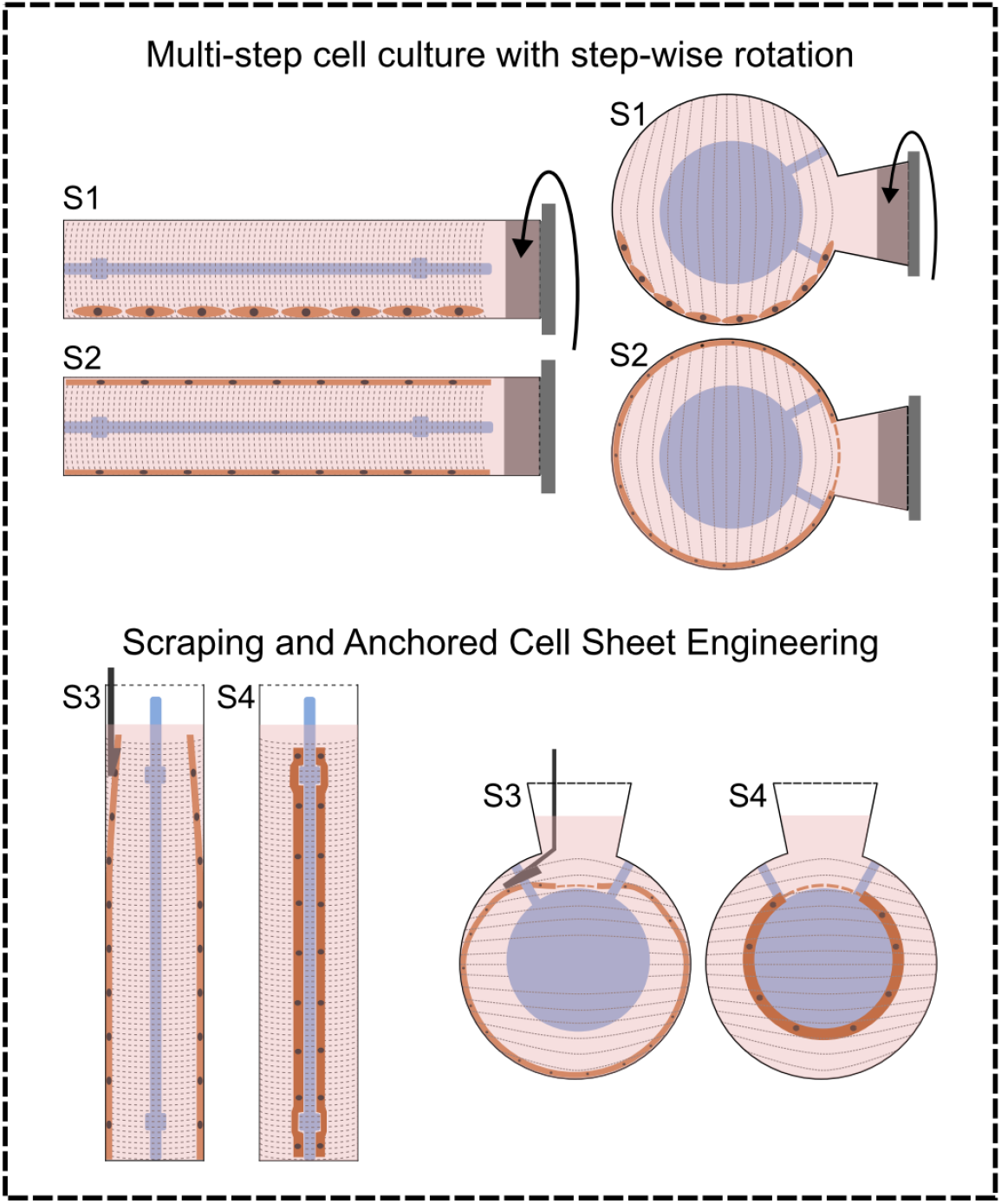

## 1. Introduction

The global demand for functional tissues and organs continues to rise, while donor availability remains limited and transplantation is burdened by the requirement for lifelong immunosuppression. These challenges have placed tissue engineering and regenerative medicine at the forefront of biomedical innovation, with the promise of patient-specific, immunologically favorable therapies [1-3]. Yet despite decades of progress, clinical translation has remained restricted, underscoring the need for biofabrication strategies that are both scalable and biomimetic [4]. Multiple approaches have been pursued to overcome this challenge. Bioprinting provides spatial control of cells and biomaterials, but struggles with resolution, cell viability during printing, and the recreation of hierarchical tissue and vascular networks [5, 6]. Xenotransplantation, though bolstered by advances in genetic engineering, continues to face immune incompatibility, zoonotic transmission risks, and ethical barriers [7]. Decellularization of native organs and then recellularizing them preserves ECM and tissue microstructure complexities but suffers from donor variability, incomplete recellularization, and limitations in scalability [8, 9]. Organoids have shown remarkable self-organization, yet their lack of vascularization and size constraints hinder scale-up to therapeutic constructs [10]. Similarly, organ-on-chip platforms yield valuable *in vitro* insights but were not designed as regenerative therapies [11]. Together, these limitations illustrate the difficulty of producing functional, clinically relevant multi-tissue constructs with existing methods.

Scaffold-free biofabrication has emerged as a promising alternative, eliminating exogenous biomaterials that can interfere with critical cell-cell and cell-matrix interactions. By enabling cells to self-organize and deposit their own ECM, scaffold-free systems generate tissues that more closely reflect *in vivo* architecture and remodeling potential. Importantly, these constructs can naturally fuse, providing a path toward modular assembly of larger, organ-like systems within the framework of bioassembloids, where functional tissue units act as standardized building blocks [12-14]. Among scaffold-free methods, spheroid-based systems have gained traction for enhancing cell-cell interactions and endogenous ECM deposition, but their growth is constrained by diffusion limits that result in necrotic cores, and their geometry is poorly suited for anisotropic tissues [15, 16]. Cell sheet engineering, pioneered with temperature-responsive surfaces, enables the harvest of intact, ECM-rich layers without enzymatic damage. Although clinical success has been demonstrated in applications such as corneal reconstruction, cardiac repair and esophageal regeneration, conventional cell sheets are fragile and restricted to planar forms, with limited maturation and ECM alignment [17-19].

Anchored Cell Sheet Engineering advanced this paradigm by introducing controlled anchorage points that reshape confluent layers after formation. These anchors create tension gradients that promote cytoskeletal realignment, ECM remodeling, and improved maturation, enabling the generation of robust sheets and fibers beyond the fragility and morphological limitations of traditional cell sheets. This strategy illustrated how proper mechanical cues could expand the repertoire of scaffold-free tissues to new form factors with enhanced mechanical and biological functionality [20]. Despite these advances, fabricating seamless three-dimensional (3D) hollow constructs remains an unmet need. Hollow architectures are fundamental to vital organs, including tubular structures such as blood vessels, trachea, bronchi, and intestine, and spherical cavities such as lung alveoli, heart chambers, and stomach. Scaffold-free replication of these geometries is essential for recapitulating native function, particularly where lumenal structures enable transport, gas exchange, or fluid containment [21, 22].

Traditional hollow tissue fabrication relies on sacrificial scaffolds or complex molding, introducing foreign materials and compromising scalability [23, 24]. Template guided assembly has been used with cell sheets wrapped around mandrels or spheroids placed on parallel shafts for making tubular constructs, but such approaches are time-consuming, produce weak seam points, challenge standardization, require sophisticated equipment, and risk disintegration under physiological conditions [25-28]. Therefore, a critical need exists for methods generating seamless, scaffold-free hollow constructs that enable direct formation of such form factors for modular bioassembly.

In this study, the Anchored Cell Sheet Engineering concept is extended to overcome this critical barrier, introducing a method for the single-step fabrication of scaffold-free hollow tubular and spherical constructs. By adapting cell culture to custom devices with curved surfaces, confluent cell-ECM layers are formed and guided to delaminate and remodel around central cores, creating robust, mechanically stable hollow tissues. The platform allows modulation of wall thickness, supports multi-layered architectures through sequential seeding of distinct cell types, and produces constructs capable of maintaining stability under fluid flow. This advancement expands the palette of scaffold-free biofabrication beyond fibers and sheets, establishing hollow structures as standardized building blocks within a bioassembloid framework for assembling complex, multi-tissue organ-like systems for regenerative medicine applications.

## 2. Results

The culture devices used in this study consisted of hollow cylinders or spheres, each containing a central shaft or sphere, respectively, to serve as an anchoring core (**Figure 1a**). These devices were fabricated by casting polydimethylsiloxane (PDMS) onto sacrificial master molds with negative designs of the intended culture devices that were 3D printed using a water-soluble polyvinyl alcohol filament (**Figure 1b**). Once the PDMS cured, the molds were dissolved with deionized water to yield the final culture devices. The complete culture system included a silicone cap with a tubing inlet for cell seeding and media exchange, a sealing wire to secure the cap in place, and a 3D printed external support to allow for manual rotation, essential for ensuring uniform cell coverage on all internal surfaces of the device (**Figure 1c**).

**Figure 1.**
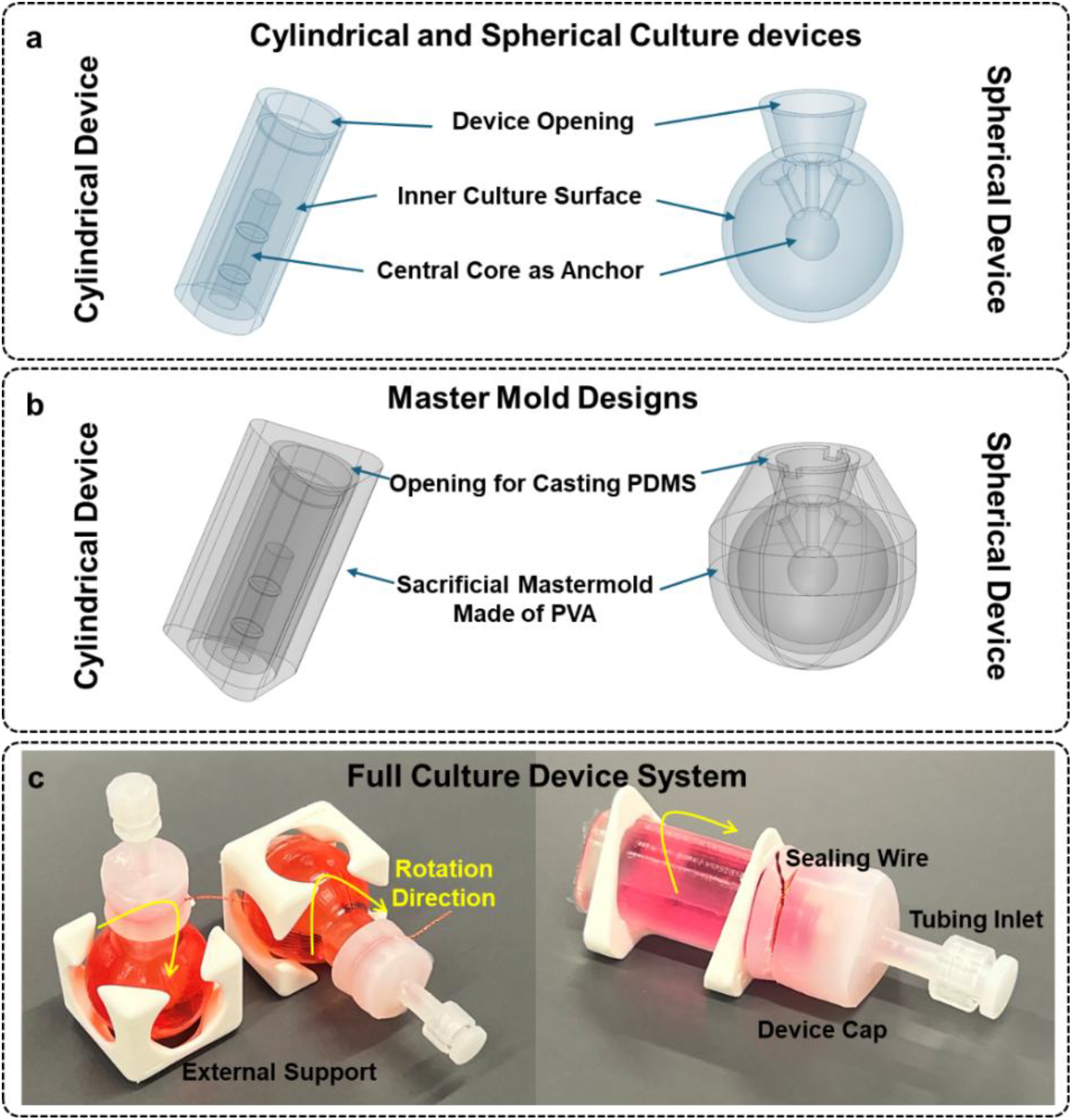
**a)** Schematics of cylindrical and spherical culture devices illustrating core anchors, device openings, and internal culture surfaces used for 2D cell culture on curved surfaces; **b)** Representative designs of master molds fabricated by 3D printing with water-soluble PVA, utilized as sacrificial templates for casting PDMS; **c)** Fully assembled culture systems showing PDMS-based cylindrical and spherical devices with Ecoflex caps and sealing wires. External supports enable controlled rotation of the culture devices following each cell addition to create a full coverage of the surface. Three rounds of cell addition in the cylindrical device require three rotations using a triangular support, while five rounds of cell addition in the spherical device require six rotations using a cubic support, as indicated by arrows.

Before cell seeding, the devices were pretreated with tannic acid to adjust hydrophilicity and promote cell adhesion. This was followed by washing, autoclaving for sterility, and coating with vitronectin for long-term cell attachment. The cell culture process (**Figure 2a)** began on day one with the introduction of 5×10^5^ primary skeletal muscle myoblasts (SkMb) in growth medium. For uniform and complete cell distribution, the devices were rotated, and the seeding process was repeated every two hours (a total of 3 and 5 times for cylindrical and spherical culture devices). On day three, the growth medium was replaced with a differentiation medium to induce myotube formation, and this medium was refreshed daily. Additional rounds of cell seeding were performed on days 4, 7, and 10. The key cell sheet engineering step was initiated on day 12 when the cell and ECM layers were gently detached from the device walls using a 1mL pipette tips and allowed to remodel overnight around the central anchoring cores (**Figure 2b**). After a total of 18 days in culture, the process yielded robust constructs that could be extracted from the cores and exhibited sufficient mechanical integrity to withstand gas and air flow (**Supplementary Videos 1 and 2**).

**Figure 2.**
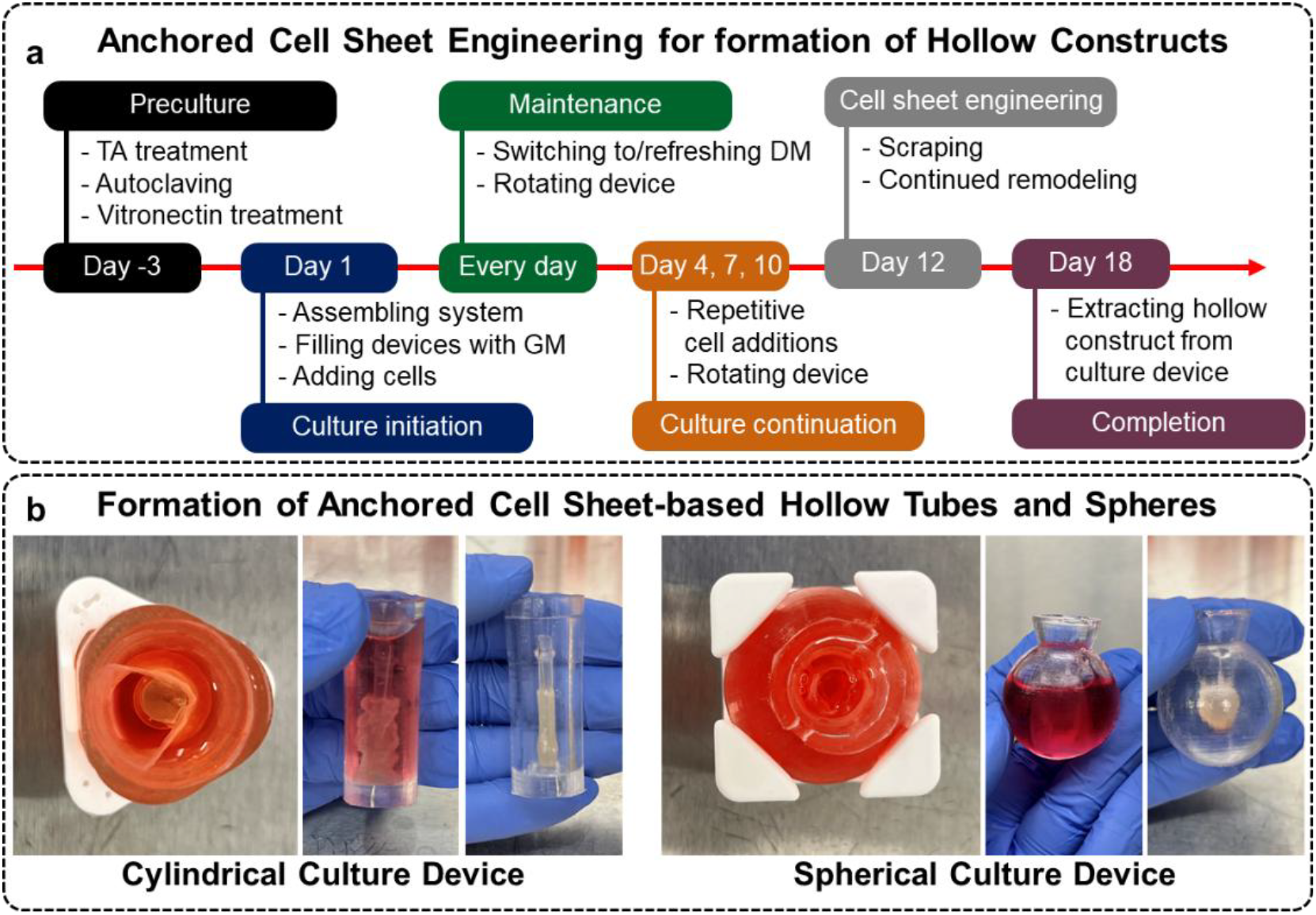
**a)** Sequential steps involved in the formation of hollow constructs using the Anchored Cell Sheet Engineering approach, beginning with preparation of the culture devices, followed by a 2D cell culture phase on curved surfaces, and concluding with cell sheet formation and anchor-mediated remodeling around the central cores. TA: tannic acid; GM: growth medium; DM: differentiation medium; **b)** Delamination of cell sheets from the PDMS surface after initial scraping results in the formation of loose cell sheets, which subsequently recognize the central cores as anchors and undergo further remodeling to generate mechanically stable constructs in a single continuous process.

The versatility of this biofabrication platform was demonstrated by creating hollow constructs of varying shapes and sizes simply by modifying the central core’s geometry (**Figures 3a and 3b**). While the outer dimensions of the culture devices remained consistent, changing the core size allowed for the fabrication of constructs with different inner diameters (**Figure 3a**). The cell sheets also displayed a remarkable capacity to remodel around more complex core shapes, such as multi-sphere and tubulo-spherical configurations, even when cultured in a simple spherical device (**Figure 3b, Supplementary Video 3**). The wall thickness of the constructs was measured using a semi-automated workflow that involved manual detection of sample areas in QuPath followed by image analysis using a custom Python script. (**Figures 3c and 4**). Quantifying H&E-stained images of the hollow tubes confirmed that wall thickness was tunable. For example, using a smaller central core (4mm ID vs. 6mm ID) in the cylindrical devices increased the wall thickness from 78.35±18.30 to 176.16±59.63 µm. Wall thickness could also be adjusted by changing the number of cell layers. Increasing the number of cell layers from four to six resulted in a wall thickness increase from 78.35±18.30 to 159.36±34.04 µm for a large core. Finally, to mimic the more complex native tissue architectures, constructs with multiple cell types were biofabricated. Sequentially layering three muscle cell layers followed by three endothelial cell layers produced a bilayered structure that was preserved during the cell sheet formation and remodelling. Distribution of the endothelial cells within these bilayred constructs that resembles the microarchitecture of blood vessels was confirmed by immunohistochemistry for the endothelial marker CD31 (**Figure 3d**).

**Figure 3.**
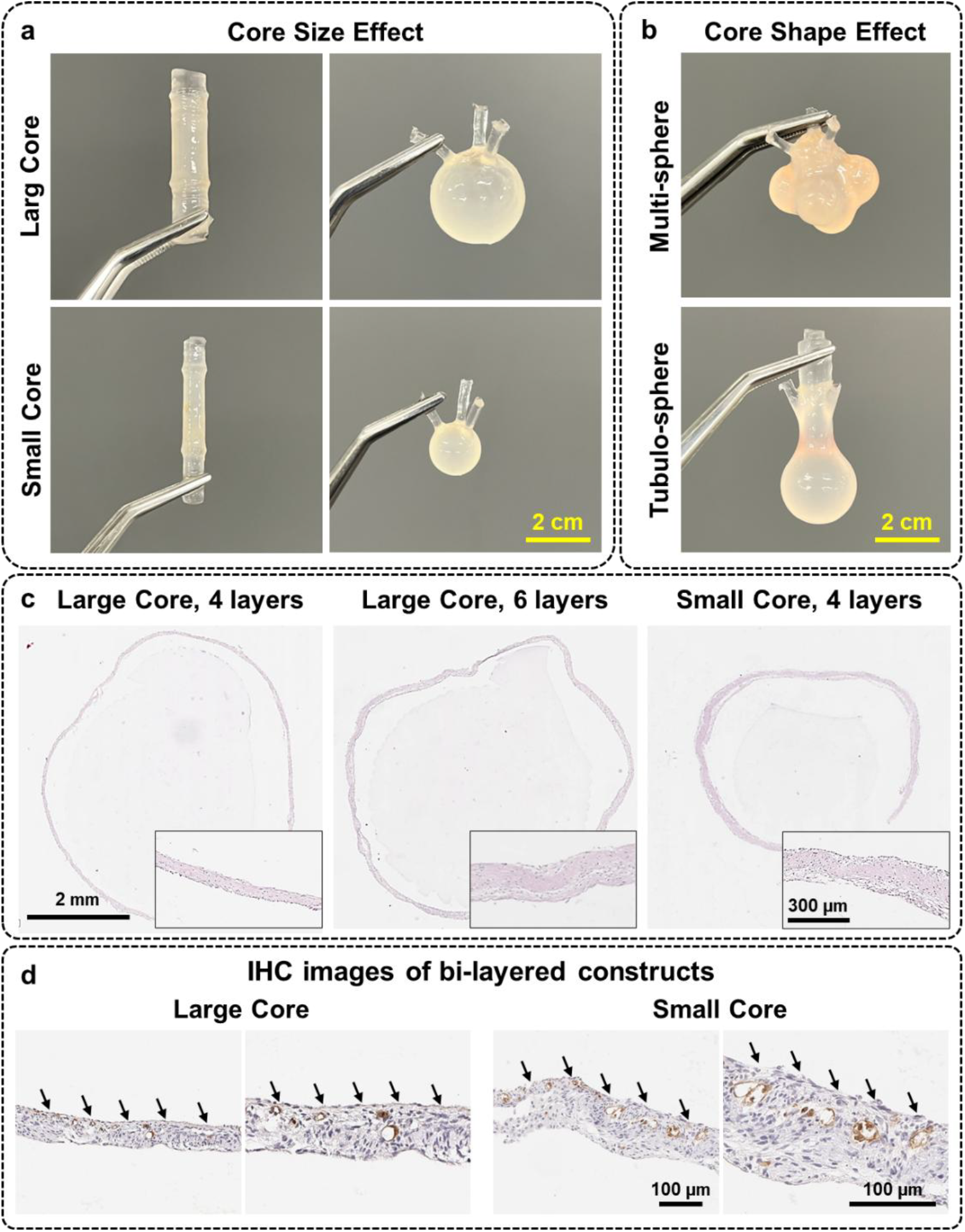
**a)** Effect of central core shape and size on the geometry of biofabricated hollow constructs. Constructs with varying inner diameters can be produced by altering the size of the core while using culture devices of consistent dimensions; **b)** Demonstration of the remodeling capacity of cell sheets to conform to complex central core geometries, including multi-sphere and tubulo-spherical shapes, even when the 2D culture is conducted on simpler spherical surfaces; **c)** Histological analysis (H&E) of hollow constructs showing modulation of wall thickness. Wall thickness can be adjusted by changing the size of the central core and/or by varying the number of cell layers grown during the 2D culture phase; **d)** Immunohistochemical characterization (CD31) of bilayered constructs formed by sequential culture of skeletal muscle and endothelial cells (3 layers each) in cylindrical culture devices, resulting in architectures that resemble physiological tissue structures such as blood vessels. Arrows indicate the area positive for endothelial marker (CD31+) in each sample. Even after remodelling, cells can maintain their initial relative positioning.

## 3. Discussion

The Anchored Cell Sheet Engineering platform has proven effective in creating highly *in vivo*-like constructs with relevant form factors, including fibers and sheets for both *in vitro* modeling and *in vivo* regenerative medicine applications [20, 29, 30]. However, the successful biofabrication of seamless hollow tubular and spherical constructs using the same concept represents a significant advance in scaffold-free tissue engineering. The results presented here show that planar 2D surfaces can be replaced with curved ones to guide cells into self-organizing into complex, seamless 3D architectures without requiring external scaffolds or manual assembly steps (**Figures 1 and 2**). This advancement directly addresses critical limitations that have constrained the field’s ability to create physiologically relevant hollow structures essential for organ engineering.

Unlike approaches such as bioprinting that struggle with cell viability and require specialized bioinks that compromise cellular functionality [31], the method introduced here preserves cellular and tissue integrity. It enables cells to deposit and organize their native ECM, resulting in superior biological relevance and mechanical properties that naturally match tissue requirements [20]. Premade scaffolds seeded with cells have also been used for these purposes. For example, Humacyte’s FDA-approved product SYMVESS™, while acellular, is made using a scaffold-based approach that requires 8-10 weeks for complete scaffold biodegradation and ECM replacement [32-34]. In contrast, the constructs here were fully formed within 18 days without any foreign biomaterial incorporation. This significant reduction in production time, combined with the elimination of scaffold-related complications, positions this technology favorably for clinical translation. Furthermore, while Humacyte’s approach is optimized for vascular applications, this platform’s versatility enables the fabrication of diverse and complex hollow geometries relevant to multiple organ systems.

The seamless nature of these constructs represents a critical advancement over traditional cell sheet wrapping methods. Previous work demonstrated tubular construct formation through manual wrapping of cell sheets around a mandrel [25, 26], but these processes are lengthy, require specialized equipment, and the formed structures could potentially exhibit mechanical failure at fusion interfaces under physiological pressures. The single-step process introduced here eliminates these weak points entirely, as evidenced by the constructs’ ability to withstand sustained gas and liquid flow without structural compromise (**Supplementary Videos 1 and 2**). The tension-mediated remodeling driven by strategic anchor placement mimics developmental morphogenetic processes where mechanical forces guide tissue organization. It’s plausible that as the cell sheet compacts around the anchor, cell-generated tension activates mechanosensitive signaling pathways. This likely involves the nuclear translocation of transcriptional co-activators like YAP and TAZ, which are known to convert mechanical cues into biological responses [35]. This could increased production and alignment of ECM components, leading to the enhanced tissue maturity and mechanical stability observed in the constructs [36]. This biomimetic approach was previously shown to result in anisotropic cell and ECM alignment similar to that of native tissues [20].

The ability to include multiple cell types and tune wall thickness (**Figures 3 and 4**) provides unprecedented control over the biofabrication of highly *in vivo*-like and physiologically relevant constructs, addressing a major limitation of existing approaches where construct properties are largely predetermined by scaffold or bioink characteristics. The demonstration of complex core geometries (**Figure 2**), including multi-sphere and tubulo-sphere configurations, extends beyond what current technologies can achieve. While organoid systems demonstrate remarkable self-organization capabilities, they remain limited to microscale structures due to diffusion constraints [10]. This approach bridges this gap by enabling fabrication of millimeter to centimeter-scale hollow constructs that could serve as functional tissue units or conduits for larger organ systems.

**Figure 4.**
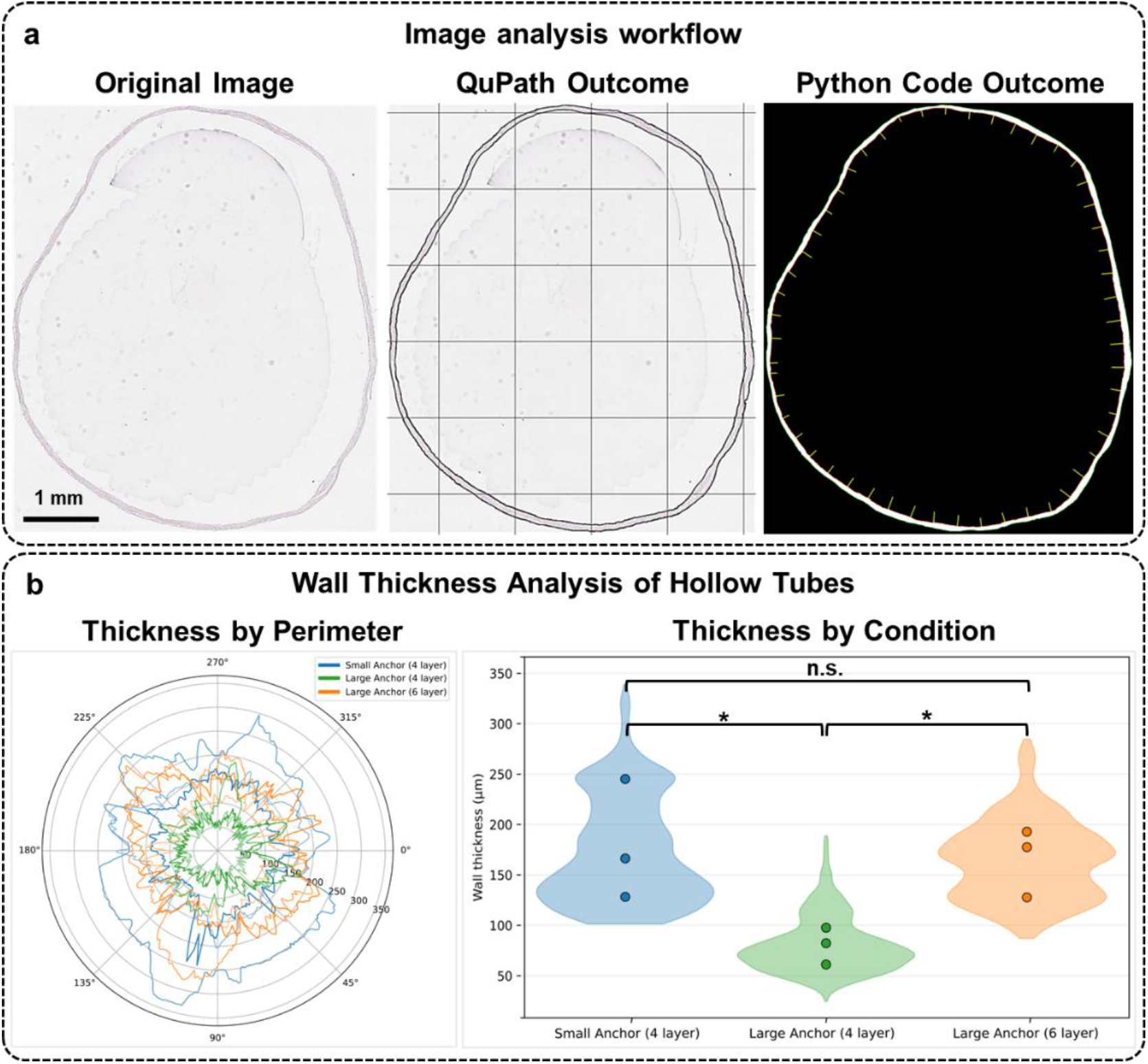
**a)** Semi-automated image analysis workflow. Regions of interest (ROI) in whole-slide images (WSI) were manually selected in QuPath (Left). The ROI contour and 1000-µm reference grid were overlaid on the original image and exported as SVG files (Center). A custom Python script was then used to generate masked ROI images and perform wall thickness measurements (Right); **b)** Wall thickness analysis of biofabricated hollow tubes. Left: polar plots showing wall thickness as a function of perimeter angle for all samples within each condition. Right: violin plots showing thickness distributions (pooled per-degree measurements) with overlaid dots representing per-sample means. Increasing number of cell layers and using a smaller core size resulted in greater wall thicknesses. n equals 3. Statistical comparisons were performed on per-sample mean values. n.s.: not significant; *: p-value < 0.05.

The Anchored Cell Sheet Engineering platform now offers a set of standardized building blocks encompassing sheets, fibers, spheroids, hollow tubes, and hollow spheres that establishes a comprehensive toolkit for bottom-up organ assembly following modular tissue engineering techniques such as bioassembloids. This modular approach contrasts sharply with monolithic organ engineering strategies that attempt to recreate entire organs in single steps. The standardized nature of these building blocks enables quality control at the component level before assembly, potentially improving overall success rates for complex tissue engineering applications by eliminating limitations such as necrotic cores and long culture times required to remodel exogenous biomaterials used as scaffolding or bioink [37, 38]. The natural fusion capacity of these scaffold-free tissue units provides a clear path for assembling these building blocks into larger structures. While recent work suggests that scaffold-free tissues can achieve seamless integration within 24-48 hours under appropriate conditions [12, 39-41], the fusion kinetics between the tissue units biofabricated in the current study need to be further evaluated.

This work establishes a versatile platform for creating scaffold-free hollow constructs to address fundamental limitations in tissue engineering, but further work is needed before it can be clinical-ready to ultimately address the critical shortage of organs for transplantation. While larger blood vessels can be fabricated using hollow tubes, broader vascularization remains a critical challenge for larger tissue constructs after bioassembly. Integration of hollow tubes with surrounding parenchymal tissue will also require sophisticated approaches. Future work should explore pre-vascularization strategies within building blocks before or after assembly. Scale-up considerations for clinical applications including automation of the process in a closed-loop manner need to be considered. Similarly, systematic, automated, and closed-loop assembly processes need to be established. Lastly, long-term stability, remodeling capacity, and function under physiological conditions require extensive evaluation to understand how these constructs adapt to chronic mechanical loading, inflammatory responses, and integration with host tissues.

## 4. Methods

### Device Fabrication

Culture devices were fabricated using polydimethylsiloxane (PDMS; Sylgard 184) cast onto master molds containing negative designs of the intended culture geometry. Master molds were fabricated using fused deposition modeling (FDM) 3D printing with water-soluble polyvinyl alcohol filament (Polymaker PolyDissolve S1 PVA). Following PDMS curing at room temperature overnight, master molds were dissolved by thorough washing with deionized water to yield the final cylindrical or spherical PDMS-based culture devices. The cylindrical devices featured an internal diameter of 14 mm and height of 45 mm, while spherical devices had an internal diameter of 30 mm. Central anchoring cores of varying dimensions were integrated into each device design (4 and 6 mm shafts and 10 and 14 mm spheres for cylindrical and spherical devices, respectively) to enable tunable construct geometries. Culture device caps were fabricated from another silicone-based resin (Ecoflex™ 00-30) and incorporated inlet tubing designed to accommodate 18-gauge needles for cell seeding and medium exchange. Sealing wires were included to secure connections between caps and culture devices. External supports made with polylactic acid (PLA) fabricated by 3D printing enabled manual rotation of culture devices during cell seeding procedures.

### Cell Culture and Cell Sheet Engineering

Culture devices were prepared for cell seeding by treating internal surfaces with aqueous tannic acid solution (Sigma-Aldrich, 403040, 50 mg/mL) for three days to enhance PDMS surface hydrophilicity. Following extensive washing with deionized water and autoclaving for sterilization, surfaces were treated with vitronectin (Gibco™, A31804, Recombinant Human Protein, Truncated, 10 µg/mL) in PBS for 1 hour at room temperature to promote cell attachment. Primary skeletal muscle myoblasts (SkMb, Sigma Aldrich, RB150) were cultured in their growth medium (Sigma Aldrich, RB151). Cell culture was initiated on day 1 by introducing 5×10^5^ cells along with growth medium to fill culture chambers (5 mL for cylindrical devices, 12 mL for spherical devices). Cell seeding was repeated every two hours with device rotation (two additional rotations for cylindrical, four for spherical devices) to ensure uniform cell distribution across internal surfaces. On day 3, growth medium was replaced with differentiation medium (Sigma Aldrich, 151D) to induce myotube formation. This medium was refreshed daily thereafter. Additional cell seeding rounds were performed on days 4, 7, and 10 using identical procedures. On day 12, devices were uncapped and cell-ECM layers were gently detached from culture surfaces using 1 mL pipette tips. Devices were recapped and incubated overnight to facilitate cell sheet remodeling and anchoring onto central cores. Cultures were maintained until day 18 to enable completion of remodeling and formation of mechanically stable constructs capable of detachment from anchoring cores.

### Histological and Immunohistochemical Analysis

Constructs were fixed in 2 wt/V% paraformaldehyde (Sigma Aldrich, 158127) in PBS for 10 minutes at room temperature, followed by dehydration through graded ethanol series and embedding in paraffin. Serial sections (5 µm thick) were cut using a microtome and mounted on glass slides. Hematoxylin and eosin (H&E) staining was performed according to standard protocols for general morphological assessment and wall thickness measurements. For immunohistochemistry (IHC) targeting CD31, tissue sections were initially deparaffinized with xylene and rehydrated through a graded ethanol series of decreasing concentrations. Antigen retrieval was carried out using citrate buffer (pH 6.0) in a pressure cooker for 20 minutes, followed by incubation with 0.3% hydrogen peroxide for 10 minutes to block endogenous peroxidase activity. The sections were then blocked with 10% donkey serum in PBS for 1 hour at room temperature. Primary antibody (Abcam, ab182981) incubation was conducted overnight at 4°C in a humidified chamber. After washing with PBS, the sections were incubated with a peroxidase-conjugated donkey anti-mouse secondary antibody (Jackson ImmunoResearch, 715-035-150). Hematoxylin was used for counterstaining, after which the sections were dehydrated, cleared, and mounted using a permanent mounting medium. All stained slides were scanned digitally at 40× magnification using a Leica Aperio AT2 scanner (Leica Biosystems, Buffalo Grove, IL) and the results were saved as whole-slide images (WSI).

### Image Analysis and Quantification

WSIs were opened in QuPath, where the hollow tubes’ cross-sections were defined as regions of interest (ROI) by manually drawing a closed contour. For each condition, three biological samples were analyzed, with two histological slides taken from different locations of each sample. A reference grid with 1000-µm spacing was overlaid to standardize spatial measurements. For each sample, the underlying image, the contour, and the reference grid were exported together as an SVG file to preserve vector geometry. Analysis proceeded using a custom Python script that isolated the ROI and converted it into a binary annular mask, which, when applied to the base image, yielded a ring-only ROI. Wall thickness was quantified using a normal-based approach. Inner and outer boundaries were extracted from the cleaned mask, and the inner boundary was resampled uniformly by arc length. At each sample point, the local outward unit normal was computed from the boundary tangent, and thickness was measured by casting a short ray along this normal. Distances were converted to micrometers using the grid-derived scale. Results were visualized as violin plots of the pooled thickness distribution for each condition, and as a combined polar overlay of all samples. For statistical analysis, mean wall thicknesses from each sample were compared across conditions. One-way ANOVA was performed to detect overall differences among groups, followed by Tukey’s honestly significant difference (HSD) post-hoc tests for all pairwise comparisons. Adjusted p-values from Tukey’s test were used to determine statistical significance, with a threshold of p-value smaller than 0.05.

## 5. Conclusions

This study establishes Anchored Cell Sheet Engineering as a versatile platform for single-step biofabrication of seamless hollow tubular and spherical constructs. By replacing planar substrates with custom curved geometries and using centrally placed cores as guiding anchors, cells form confluent inner linings that remodel via controlled delamination into mechanically robust, biologically relevant 3D architectures. Dimensions and shapes are tunable through core design, and multiple cell types can be layered to produce biomimetic, multi-layer structures with native ECM deposition and anisotropic alignment. Together with sheets, fibers, and spheroids, these hollow forms constitute standardized scaffold-free building blocks that underpin the bioassembloid concept for bottom-up assembly of larger tissues and, ultimately, organ-scale constructs. This approach removes weak seams typical of template-guided methods and avoids the prolonged remodeling, specialized biomaterials, and viability losses common to scaffold-based and bioprinting strategies, enabling stable hollow geometries within days. Priorities ahead include demonstrating reproducible, autonomous fusion among modules and with host tissues *in vivo*, integrating pre-vascularization and perfusion-friendly design rules, establishing quantitative potency and functional metrics, and automating manufacturing for clinical scale and regulatory compliance. By uniting modularity, scalability, and physiological relevance, Anchored Cell Sheet Engineering offers a practical, biomimetic route to scaffold-free organ engineering.

## Acknowledgment

The authors expressed gratitude to Caroline O’Neil from Robarts Research Institute, Western University, London, Canada for performing the histology assessment. All other experiments were performed at Velocity Incubator, Waterloo, Canada. The authors appreciate the support from Velocity Incubator’s staff.

## Data Availability Statement

The custom Python script used in analyzing the data is available on GitHub at https://github.com/Evolved-Bio/TissueEngineeringGrail. The STL files generated and used in this study have been deposited in the Zenodo repository and are accessible at https://doi.org/10.5281/zenodo.17085955. Any other data used in the study but not included here is available upon request.

### Declaration of use of generative AI

During the preparation of this work, the authors used Open AI’s ChatGPT and Anthropic’s Claude in order to grammatically edit the manuscript and debug the Python scripts. After using these tools, the authors reviewed and edited the content as needed and take full responsibility for the content accuracy.

## References

1. Giwa, S., et al., The promise of organ and tissue preservation to transform medicine. Nat Biotechnol, 2017. 35(6): p. 530–542.

2. Jain, A. and R. Bansal, Applications of regenerative medicine in organ transplantation. J Pharm Bioallied Sci, 2015. 7(3): p. 188–94.

3. Peloso, A., et al., Current achievements and future perspectives in whole-organ bioengineering. Stem Cell Res Ther, 2015. 6(1): p. 107.

4. O’Donnell, B.T., et al., Beyond the Present Constraints That Prevent a Wide Spread of Tissue Engineering and Regenerative Medicine Approaches. Front Bioeng Biotechnol, 2019. 7: p. 95.

5. Ozbolat, I.T. and Y. Yu, Bioprinting toward organ fabrication: challenges and future trends. IEEE Trans Biomed Eng, 2013. 60(3): p. 691–9.

6. Huang, G., et al., Applications, advancements, and challenges of 3D bioprinting in organ transplantation. Biomater Sci, 2024. 12(6): p. 1425–1448.

7. Eisenson, D.L., Y. Hisadome, and K. Yamada, Progress in Xenotransplantation: Immunologic Barriers, Advances in Gene Editing, and Successful Tolerance Induction Strategies in Pig-To-Primate Transplantation. Front Immunol, 2022. 13: p. 899657.

8. Scarritt, M.E., N.C. Pashos, and B.A. Bunnell, A review of cellularization strategies for tissue engineering of whole organs. Front Bioeng Biotechnol, 2015. 3: p. 43.

9. Hillebrandt, K.H., et al., Strategies based on organ decellularization and recellularization. Transpl Int, 2019. 32(6): p. 571–585.

10. Takebe, T. and J.M. Wells, Organoids by design. Science, 2019. 364(6444): p. 956–959.

11. Ma, C., et al., Organ-on-a-Chip: A New Paradigm for Drug Development. Trends Pharmacol Sci, 2021. 42(2): p. 119–133.

12. De Pieri, A., Y. Rochev, and D.I. Zeugolis, Scaffold-free cell-based tissue engineering therapies: advances, shortfalls and forecast. NPJ Regen Med, 2021. 6(1): p. 18.

13. Liu, K.C., et al., Scaffold-free 3D culture systems for stem cell-based tissue regeneration. APL Bioeng, 2024. 8(4): p. 041501.

14. Kurniawan, N.A., The ins and outs of engineering functional tissues and organs: evaluating the in-vitro and in-situ processes. Curr Opin Organ Transplant, 2019. 24(5): p. 590–597.

15. Mironov, V., et al., Organ printing: tissue spheroids as building blocks. Biomaterials, 2009. 30(12): p. 2164–74.

16. Baptista, L.S., et al., Bioprinting Using Organ Building Blocks: Spheroids, Organoids, and Assembloids. Tissue Eng Part A, 2024. 30(13-14): p. 377–386.

17. Zurina, I.M., et al., Towards clinical translation of the cell sheet engineering: Technological aspects. Smart Materials in Medicine, 2023. 4: p. 146–159.

18. Hu, D., et al., The preparation methods and types of cell sheets engineering. Stem Cell Res Ther, 2024. 15(1): p. 326.

19. Thummarati, P., et al., Recent Advances in Cell Sheet Engineering: From Fabrication to Clinical Translation. Bioengineering (Basel), 2023. 10(2).

20. Shahin-Shamsabadi, A. and J. Cappuccitti, Anchored Cell Sheet Engineering: A Novel Scaffold-Free Platform for in vitro Modeling. Advanced Functional Materials, 2024. 34(13): p. 2308552.

21. Pien, N., et al., Tubular Bioartificial Organs: From Physiological Requirements to Fabrication Processes and Resulting Properties. A Critical Review. Cells Tissues Organs, 2022. 211(4): p. 420–446.

22. Holland, I., et al., 3D biofabrication for tubular tissue engineering. Biodes Manuf, 2018. 1(2): p. 89–100.

23. Li, S., et al., Recent advances in 3D printing sacrificial templates for fabricating engineered vasculature. MedComm – Biomaterials and Applications, 2023. 2(3): p. e46.

24. Laowpanitchakorn, P., et al., Biofabrication of engineered blood vessels for biomedical applications. Sci Technol Adv Mater, 2024. 25(1): p. 2330339.

25. L’Heureux, N., et al., A completely biological tissue-engineered human blood vessel. Faseb j, 1998. 12(1): p. 47–56.

26. L’Heureux, N., et al., Human tissue-engineered blood vessels for adult arterial revascularization. Nat Med, 2006. 12(3): p. 361–5.

27. Aguilar, I.N., et al., Scaffold-free Bioprinting of Mesenchymal Stem Cells with the Regenova Printer: Optimization of Printing Parameters. Bioprinting, 2019. 15.

28. Kim, M.H., et al., High-Throughput Bioprinting of Spheroids for Scalable Tissue Fabrication. bioRxiv, 2024.

29. Shahin-Shamsabadi, A. and J. Cappuccitti, In Vivo-Like Scaffold-Free 3D In Vitro Models of Muscular Dystrophies: The Case for Anchored Cell Sheet Engineering in Personalized Medicine. Adv Healthc Mater, 2025. 14(12): p. e2404465.

30. Shahin-Shamsabadi, A. and J. Cappuccitti, Muscle-specific acellular ECM fibers made with anchored cell sheet engineering support regeneration in rat models of volumetric muscle loss. Acta Biomater, 2025. 200: p. 416–431.

31. Kim, S.J., et al., Bioprinting Methods for Fabricating In Vitro Tubular Blood Vessel Models. Cyborg Bionic Syst, 2023. 4: p. 0043.

32. Nash, K.M., et al., Evaluation of tissue-engineered human acellular vessels as a Blalock-Taussig-Thomas shunt in a juvenile primate model. JTCVS Open, 2023. 15: p. 433–445.

33. Sokolov, O., et al., Use of bioengineered human acellular vessels to treat traumatic injuries in the Ukraine-Russia conflict. Lancet Reg Health Eur, 2023. 29: p. 100650.

34. Kirkton, R.D., et al., Evaluation of vascular repair by tissue-engineered human acellular vessels or expanded polytetrafluoroethylene grafts in a porcine model of limb ischemia and reperfusion. J Trauma Acute Care Surg, 2023. 95(2): p. 234–241.

35. Panciera, T., et al., Mechanobiology of YAP and TAZ in physiology and disease. Nat Rev Mol Cell Biol, 2017. 18(12): p. 758–770.

36. Humphrey, J.D., E.R. Dufresne, and M.A. Schwartz, Mechanotransduction and extracellular matrix homeostasis. Nat Rev Mol Cell Biol, 2014. 15(12): p. 802–12.

37. Nichol, J.W. and A. Khademhosseini, Modular Tissue Engineering: Engineering Biological Tissues from the Bottom Up. Soft Matter, 2009. 5(7): p. 1312–1319.

38. Ouyang, L., et al., Assembling Living Building Blocks to Engineer Complex Tissues. Advanced Functional Materials, 2020. 30(26): p. 1909009.

39. Li, M., et al., Cell sheet technology: a promising strategy in regenerative medicine. Cytotherapy, 2019. 21(1): p. 3–16.

40. Susienka, M.J., B.T. Wilks, and J.R. Morgan, Quantifying the kinetics and morphological changes of the fusion of spheroid building blocks. Biofabrication, 2016. 8(4): p. 045003.

41. Kosheleva, N.V., et al., Cell spheroid fusion: beyond liquid drops model. Sci Rep, 2020. 10(1): p. 12614.

